# Geometrical constraints greatly hinder formin mDia1 activity

**DOI:** 10.1101/671842

**Authors:** Emiko L. Suzuki, Bérengère Guichard, Guillaume Romet-Lemonne, Antoine Jégou

**Author notes:** Corresponding authors: Antoine Jégou and Guillaume Romet-Lemonne, Université de Paris, Institut Jacques Monod, CNRS, 15 rue Hélène Brion, F-75013 Paris, France, and.

## Abstract

Formins are one of the central players in the assembly of most actin networks in cells. The sensitivity of these processive molecular machines to mechanical tension is now well established. However, how the activity of formins is affected by geometrical constraints related to network architecture, such as filament crosslinking and formin spatial confinement, remains largely unknown. Combining microfluidics and micropatterning, we reconstituted *in vitro* mDia1 formin-elongated filament bundles induced by fascin, with different geometrical constraints on the formins, and measured the impact of these constraints on formin elongation rates and processivity. When filaments are not bundled, formins can be anchored to static or fluid surfaces, by either end of the proteins, without affecting their activity. We show that filament bundling by fascin reduces both unanchored formin elongation rate and processivity. Strikingly, when filaments elongated by surface-anchored formins are cross-linked together, formin elongation rate immediately decreases and processivity is reduced, up to 24-fold, depending on the cumulative impact of formin rotational and translational freedoms. Our results reveal an unexpected crosstalk between the constraints at the filament and the formin levels. We anticipate that in cells, the molecular details of formin anchoring to the plasma membrane, strongly modulate formin activity at actin filament barbed ends.

## Introduction

To perform complex cellular functions and mechanotransduction at the micron scale, actin filaments assemble to create networks that vary in size, structure and dynamics. Actin filaments are continuously generated, polymerized, crosslinked to each other or attached to membranous cellular compartments. The intertwining of actin filament assembly and crosslinking is tightly regulated in space and time to shape the various cytoskeletal structures, such as the cell cortex, stress fibers, or transverse arcs (Blanchoin et al., 2014).

Formins, together with Ena/VASPs, are actin binding proteins that have the unique ability to processively track actin filament barbed ends and increase their elongation rates (Hansen and Mullins, 2010; Kovar and Pollard, 2004; Romero et al., 2004; Zigmond et al., 2003). Members of the formin family are head-to-tail homodimers with two functional formin homology domains, FH1 and FH2 (recently reviewed in (Courtemanche, 2018; Zimmermann and Kovar, 2019)). The FH2 homodimer interacts with the last subunits of the actin filament barbed end, adopts different conformations in rapid equilibrium, and gates actin monomer addition or removal from the barbed end (Otomo et al., 2005). The FH1 domains are disordered domains harboring multiple polyproline tracks to which profilin and profilin-actin complexes bind. These two functional domains work together to speed up actin filament barbed end elongation (Paul and Pollard, 2008; Vavylonis et al., 2006).

The mechanosensitivity of formins is now well established. Applying a pulling force on a formin bound to an actin filament led to an increase in barbed end elongation rate for mDia1 (Jégou et al., 2013; Kubota et al., 2017; Yu et al., 2017, 2018), mDia2 (Cao et al., 2018; Zimmermann et al., 2017) and Bni1p (Courtemanche et al., 2013), while the opposite effect was observed for Cdc12 (Zimmermann et al., 2017) and for Bni1p in absence of profilin (Courtemanche et al., 2013). Recently, mDia1 and mDia2 formin processivity was shown to depend on the efficiency of FH1 domains to bind, with the help of profilin, to the actin filament barbed end and increase the lifetime of the FH2 domains interaction with the barbed end (Cao et al., 2018). Most importantly, formin mDia1 and mDia2 processivity was shown to be severely reduced when a pulling force was applied on those formins, with a 3 pN pulling force increasing the formin dissociation rate more than 10-fold (Cao et al., 2018).

These observations have provided novel insights into the molecular details of formin function and crucial evidence for the high sensitivity of formins to biochemical and mechanical conditions to which they are exposed. However, they were obtained in cases where formins elongated single isolated actin filaments. This situation is rarely encountered in cells where membrane-associated formins elongate filaments which are cross-linked together into actin networks.

Here, using *in vitro* approaches, we investigated the activity of mDia1(FH1FH2DAD) formins (hereafter referred to as formins) in various geometrical configurations, with the fascin-induced bundle geometry as a case study.

As readouts of formin activity, we monitored the elongation rate of filament barbed ends and the formin detachment rate. We first examined, separately, the consequences of geometrical constraints on formins (anchoring) and on filaments (bundling). Different anchoring conditions were tested, binding formins from either end (FH1 or FH2 side), on a solid or a fluid surface. None of these conditions had a significant impact on formin activity at the barbed ends of independent, individual filaments (figure 1). When we bundled filaments elongating with free, unanchored formins, we found that bundling on its own slowed down elongation and enhanced formin detachment (figure 2). We next combined filament bundling and formin anchoring and found that formin activity was affected further, in ways that depended on the anchoring features (figures 3-5). These results show that details of formin anchoring, which appear inconsequential when elongating single filaments, become crucial when filaments are bundled together. We determined that, in order to function efficiently when filaments are bundled, anchored formins must have both rotational freedom (figure 4) and translational freedom (figure 5).

**figure 1:**
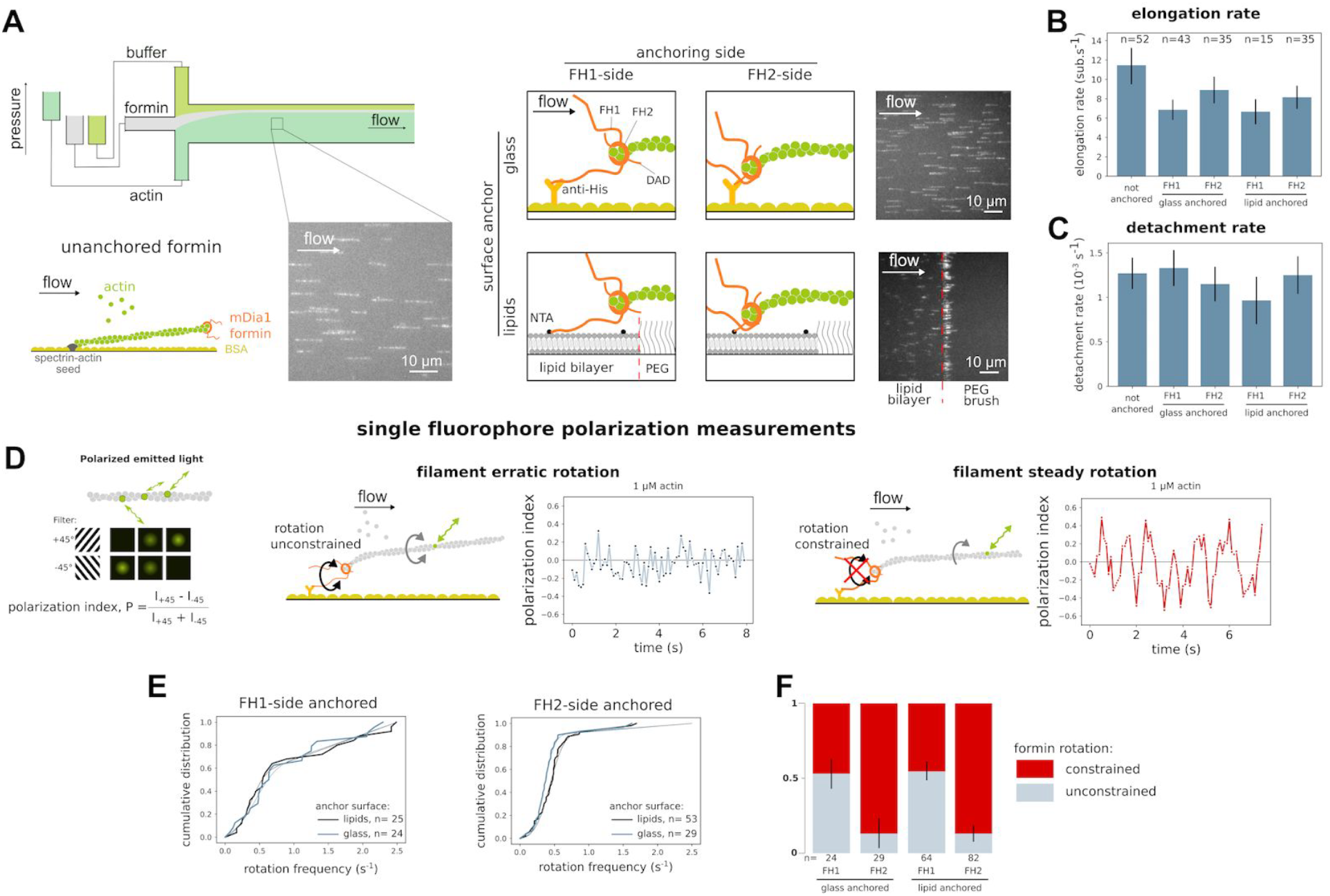
Formin anchoring may impact its rotational freedom but does not affect single actin filament elongation. (A) In a microfluidics chamber, shown in upper left, formin-induced actin filament elongation is monitored for unanchored formins (bottom left) or for formins specifically anchored to either glass or a lipid bilayer, by a 6x Histidine tag located either at their N- or C-terminus. Typical fields of view showing 15% Alexa 488-labeled actin filaments aligned by the microfluidics flow. (B) Formin-induced actin filament mean elongation rate, in the presence of 0.2 µM actin and 2 µM profilin, as a function of formin anchoring (error bar is sample standard deviation). (C) Formin detachment rate as a function of formin anchoring in the presence of 0.2 µM actin and 2 µM profilin (error bar is standard error, see Methods). (D) The polarization of the emitted light indicates the orientation of a single fluorescently-labeled actin subunit. The polarization index P = (I_+ 45_ − I_−45_)/(I_+45_ + I_−45_) is determined by measuring the emitted intensity through two orthogonal polarization filters (I_+ 45_ and I_−45_). Depending on the rotation constraints at the anchoring point, an actin filament elongated by an anchored formin will exhibit either an erratic variation of polarization (left), or a steady oscillation of the polarization signal (right). For each situation sketched, a typical experimental data curve is shown as an example. (E) Cumulative distributions of the main frequency of the polarization signal of the light emitted by a single fluorescently labelled-actin subunit, for filaments elongated by (left) FH1-anchored formins, or (right) FH2-anchored formins, anchored either on glass or lipids, in the presence of 1 µM actin. Experimental data are fitted by the weighted sum of the cumulative distribution functions of a normal distribution and a random distribution. (F) The fraction of formins identified as rotationally constrained or rotationally unconstrained were determined by fitting the cumulative distributions shown in panel E, for the four different anchoring situations shown in panel A.

**Figure 2:**
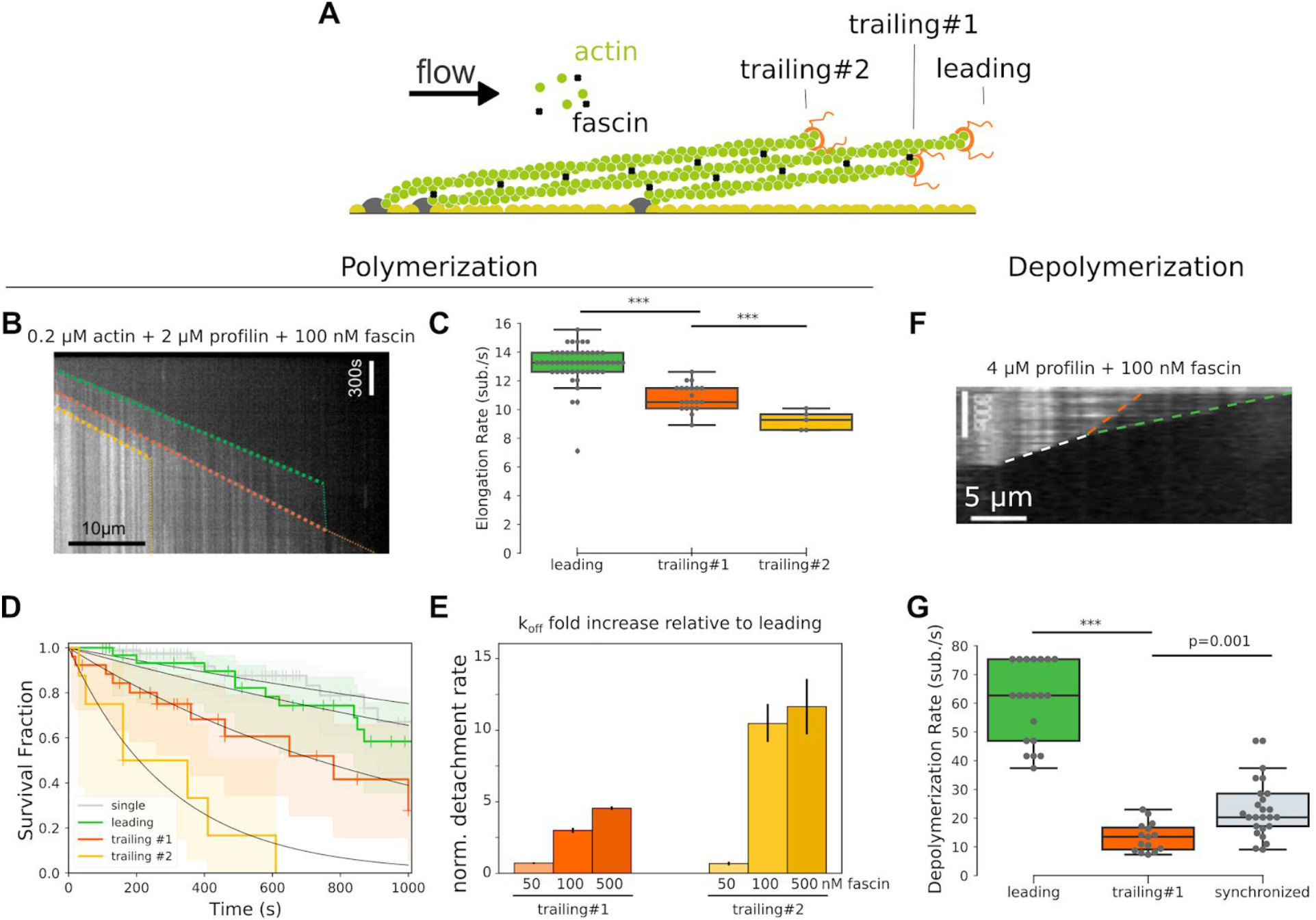
Fascin-induced bundling slows down trailing actin filament barbed end elongation and increases trailing formin detachment rate. (A) Sketch depicting a fascin-induced bundle where barbed ends are elongated by unanchored formins, identified as ‘leading’, ‘trailing #1’ and ‘trailing #2’ depending on the relative barbed end positions. (B) Kymograph showing the elongation of bundled filaments by formins, in the presence of 0.2 µM actin, 2 µM profilin and 100 nM fascin, for ‘leading’ (green dashed line), ‘trailing #1’ (orange dashed line) and ‘trailing #2’ (yellow dashed line) formins. (C) Mean filament elongation rate for the three types of unanchored formins (N= 52, 21, 5 for ‘leading’, ‘trailing #1’ and ‘trailing #2’ respectively; error bar is sample standard deviation). (D) Survival fraction of formin bound barbed ends as a function of time for single filaments (gray, n=80) or bundled filaments whose barbed end is ‘leading’ (green, n=34), ‘trailing #1’ (orange, n=28) and ‘trailing #2’ (yellow, n=8), fitted by a monoexponential decay function. (E) Normalized detachment rate of trailing formins, relative to leading formin detachment rate, as a function of fascin concentration. (error bar is standard error of the fit) (F) Kymograph showing the depolymerization of bundled filaments by formins, in the presence of 4 µM profilin and 100 nM fascin, for ‘leading’ formin (green dashed line), ‘trailing #1’ formin (orange dashed line) and ‘synchronized’ leading and trailing (white dashed line) formins. (G) Mean filament depolymerization rate for the three types of unanchored formins (N= 20, 15, 25 for ‘leading’, ‘trailing #1’ and ‘synchronized’ formins respectively; error bar is sample standard deviation).

**Figure 3:**
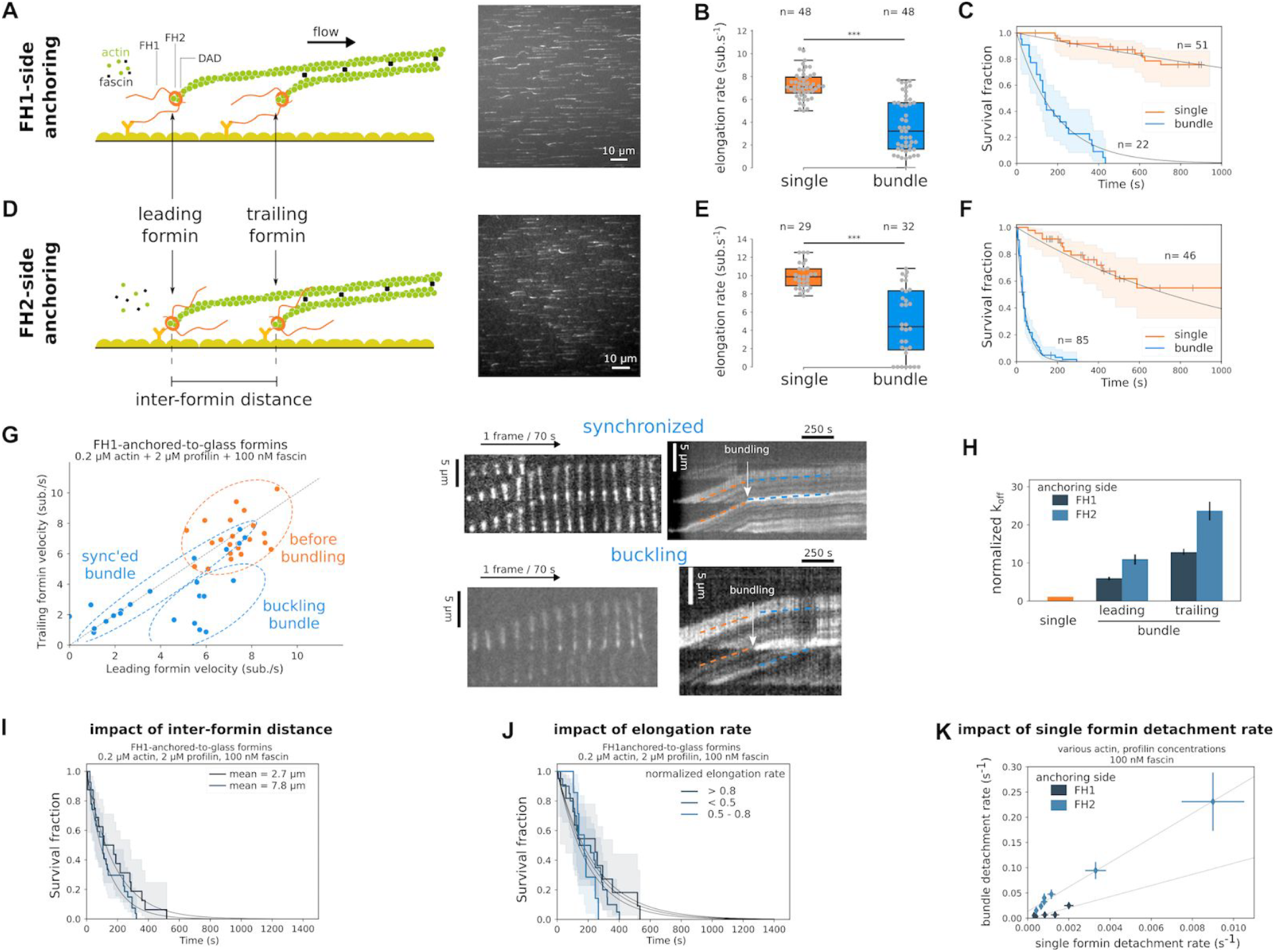
Fascin bundling affects glass-anchored formin elongation and detachment rates. (A and D) Sketches of a 2-filament bundle elongated by (A) FH1-anchored-to-glass or (D) FH2-anchored-to-glass formins, and the respective typical fields of view showing Alexa 488-labeled actin filaments bundling together. (B and E) Barbed end elongation rate distributions of single or bundled filaments elongated by (B) FH1-anchored-to-glass or (E) FH2-anchored-to-glass formins, in the presence of 0.2 µM actin, 2 µM profilin and 100 nM fascin. (C and F) Survival fractions of formin-bound single or leading+trailing filaments (referred as ‘bundle’) for (C) FH1-anchored-to-glass or (F) FH2-anchored-to-glass formins, in the presence of 0.2 µM actin, 2 µM profilin and 100 nM fascin. (G) Pairwise elongation rates for trailing and leading filaments in 2-filament bundles for FH1-anchored-to-glass formins, before or during bundle elongation, in the presence of 0.2 µM actin, 2 µM profilin and 100 nM fascin. Typical Images and corresponding kymographs of 2-filament bundles where bundling leads to either (top) ‘synchronized’ elongation of the two filaments, or (bottom) ‘buckling’ of the leading filament. (H) Normalized detachment rate of the leading and trailing FH1-anchored-to-glass or FH2-anchored-to-glass formins, relative to the detachment rate of single formins (error bars are standard error). (I) Survival fractions of FH1-anchored-to-glass formin-bound bundled filaments for two populations where the inter-formin distance is 2.7 ± 0.8 and 7.8 ± 3.3 µm (average ± standard deviation, n= 16 and 27 bundles respectively). (J) Survival fractions of FH1-anchored-to-glass formin-bound bundled filaments for three populations where the normalized elongation rate relative to the elongation rate before bundling is higher than 0.8, lower than 0.5 or in between (n= 11, 20 and 7 bundles respectively). (K) Bundle detachment rate as a function of single formin detachment rate, at 100 nM fascin, obtained by varying actin and profilin concentrations, for FH1-anchored-to-glass or FH2-anchored-to-glass formins (error bars are standard error).

**figure 4:**
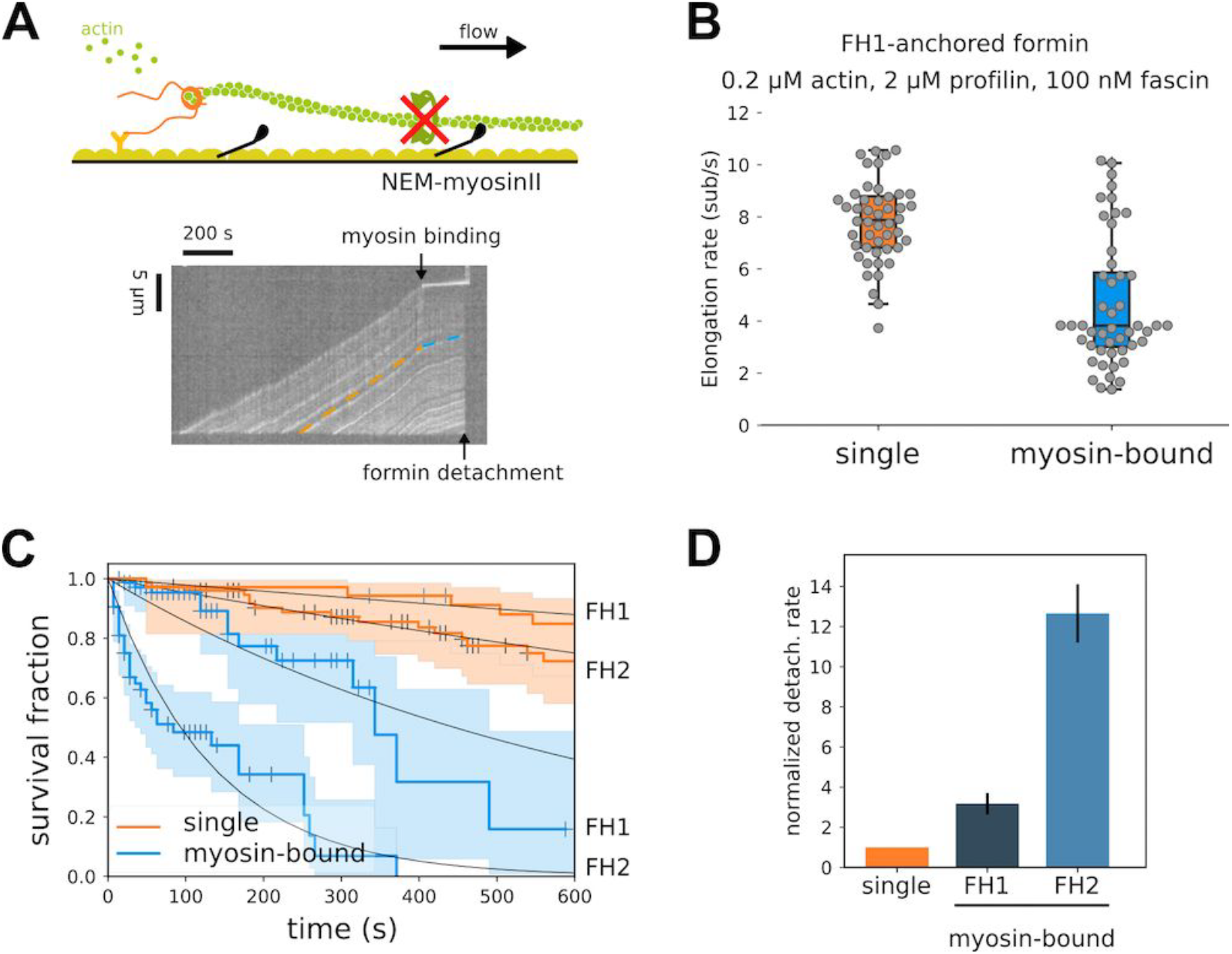
Single surface attachment reduces single formin elongation and processivity. (A) (top) Sketch of a a glass-anchored formin elongating a single filament whose rotation is blocked by surface attachment upon binding of a NEM-myosin to filament side. (bottom) Kymograph showing an actin filament elongated by a FH1-anchored-to-glass formin and whose elongation is slowed down upon filament binding to a surface-bound NEM-myosin. (B) Distribution of elongation rates of individual actin filaments, elongated by FH1-anchored-to-glass formins, bound or not to NEM-myosins attached to the glass surface (n=48 for both conditions; error bars are sample standard deviations). (C) Survival fractions of formin-bound single filaments attached to the surface via a NEM-myosin or not, for FH1-anchored-to-glass (n=78 and 35 filaments for NEM-myosin-bound or not) or FH2-anchored-to-glass formins (n=53 and 76 filaments for NEM-myosin-bound or not), in the presence of 0.2 µM actin and 2 µM profilin. (D) Normalized formin detachment rates for either FH1-anchored-to-glass or FH2-anchored-to-glass individual formins, whose elongating filaments are rotationally blocked by a surface-attached NEM-myosin, relative to glass-anchored formins elongating rotationally unconstrained single filaments.

**figure 5:**
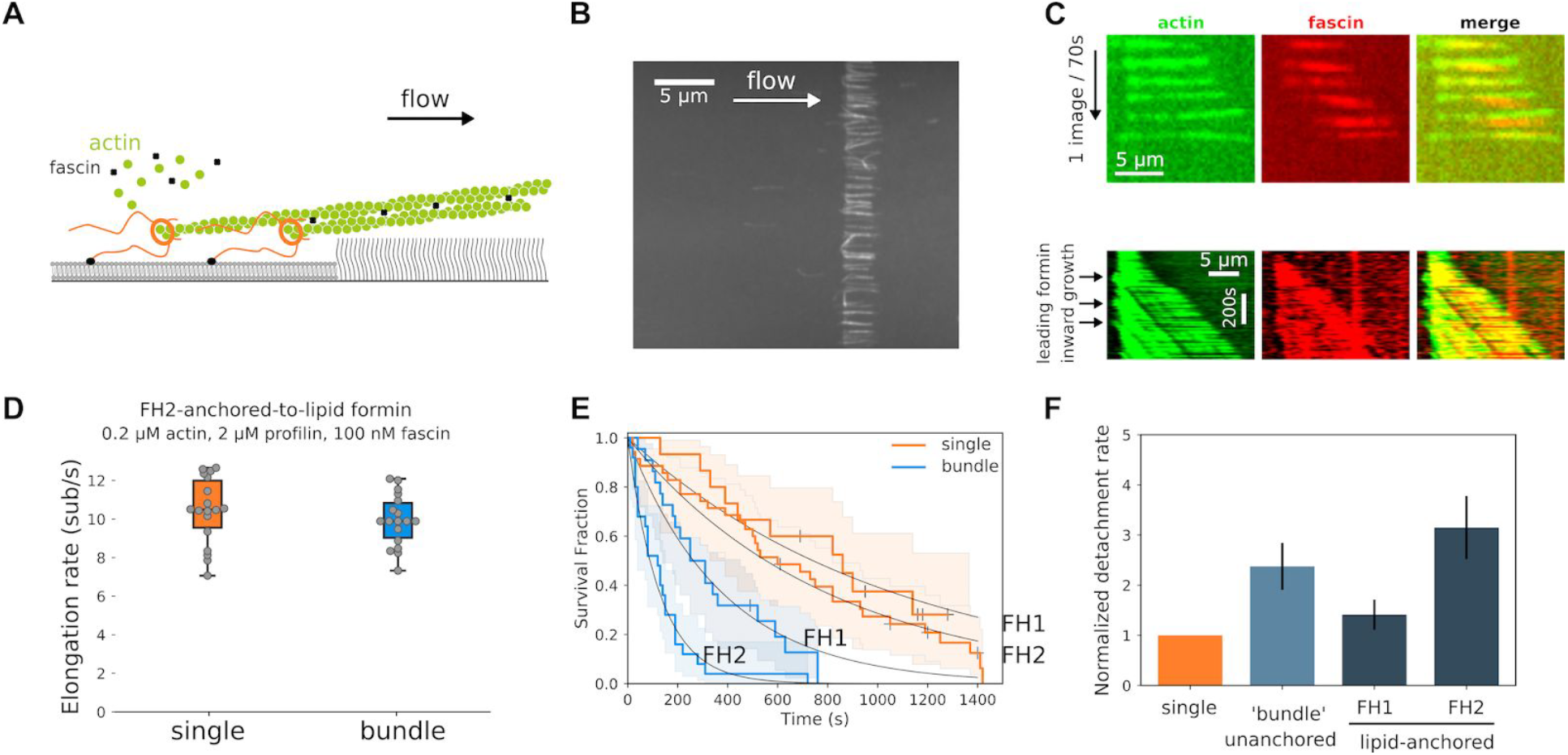
Fascin bundling moderately affects lipid-anchored formin detachment rate, but not elongation rate. (A) Sketch of a fascin-induced 2-filament bundle elongated by lipid-anchored formins in a microfluidics flow. (B) Typical field of view of filament bundles elongated by FH1-anchored-to-lipid formins at the edge of a lipid square pattern, in the presence of 0.2 µM actin, 2µM profilin and 100 nM fascin, in a microfluidics flow. (C) Raw data of a 2-filament bundles imaged with both 15% Alexa 488-labeled actin and 100% Alexa 568-labeled fascin; Bottom: Kymograph of a 2-filament bundle where a leading filament moves upstream as its elongation rate is higher than the elongation rate of the trailing filament. (D) Elongation rate distributions of single filaments (n=18) or 2-filament bundles (n=18) elongated by FH2-anchored-to-lipid formins (Student’s t-test p-value=0.34). (E) Survival fractions of lipid-anchored formin-bound single filaments or 2-filament bundles, for FH1-anchored-to-lipid (n=15 and 22 filaments, for single filaments and 2-filament bundles respectively) or FH2-anchored-to-glass formins (n=35 and 25 filaments, for single filaments and 2-filament bundles respectively), in the presence of 0.2 µM actin, 2 µM profilin and 100 nM fascin. (F) Normalized individual formin detachment rates for either FH1-anchored-to-lipids or FH2-anchored-to-lipids individual formins elongating 2-filament bundles, relative to the detachment rate of formins elongating single, unbundled filaments (orange bar). For comparison, the average detachment rate of both leading and trailing unanchored formins is shown (light blue bar). (error bars are standard errors).

## Results

### Single formin mDia1 can be attached to surfaces without affecting its activity

In cells, activated formins are bound to membranes compartments, via direct binding to lipids or through protein complexes (Chakrabarti et al., 2018; Shao et al., 2015; Skau et al., 2015). To study the impact of formin anchoring, we used an *in vitro* microfluidics approach where we tested four different anchoring schemes and compared them to situations where the formins were not anchored: formins were anchored by either their N-terminus or their C-terminus, and either to proteins adsorbed on glass (referred to as “FH1-anchored-to-glass” and “FH2-anchored-to-glass” formins) or to a freely diffusing lipid bilayer bordered by a PEG brush (referred to as “FH1-anchored-to-lipids” and “FH2-anchored-to-lipids” formins) (see figure 1A and Methods).

In all these configurations, individual formins behaved similarly in terms of elongation rate (figure 1B) and processivity (figure 1C). These results are in agreement with previous reports for glass-anchored formins (Cao et al., 2018; Jégou et al., 2013; Kubota et al., 2017). In the lipid-anchored formin configuration, the microfluidics flow dragged the filament-elongating formins to the edge of the lipid pattern, where their activity was unaffected. The PEG brush bordering the lipid pattern thus appears to differ from solid chromium barriers, which were reported to affect the activity of other formins when they were similarly dragged by a flowing solution (Courtemanche et al., 2013).

To further characterize our different anchoring schemes, we looked at how anchoring formins either via their FH1 or via their FH2 side could affect their ability to freely rotate around their surface attachment point. This aspect is important because, due to the helical structure of the actin filament, it appears that the formin needs to be able to rotate around the filament axis as it tracks the growing barbed end (Kovar and Pollard, 2004; Mizuno et al., 2011; Otomo et al., 2005). It has recently been reported that different surface-anchoring schemes, putting different constraints on formin rotation may affect its ability to elongate filaments (Yu et al., 2017). Here, for FH1-side anchored formins, the FH1 domain can be considered as a flexible linker with a contour length of ~40 nm, connecting the formin to the surface (Horan et al., 2018). In contrast, when formins are anchored by their FH2 side, the DAD domain, which lies downstream of FH2 and is composed of a short alpha-helix followed by an unstructured region, can be considered as a linker of ~10 nm in contour length (Kühn and Geyer, 2014). FH2-anchoring thus constitutes a shorter tether than FH1-anchoring, and we anticipated that it would affect the formin’s ability to rotate around its anchoring point as the tether length is similar to the FH2-bound actin filament barbed end diameter (~8-10 nm) (Otomo et al., 2005).

In order to assess filament rotation around its main axis, we measured the polarization of the light emitted by a single fluorescently labeled actin subunit incorporated within the filament (figure 1D) (Mizuno et al., 2011; Wioland et al., 2019). This allowed us to determine the polarization index of this subunit, which varies over time as the filament rotates around its axis (which itself points in the fixed direction of the flow). If the filament cannot rotate around its axis, the polarization index remains constant (supp. figure 1). If the filament is anchored to the surface via a formin, we can expect two main outcomes depending on the formin’s ability to rotate: (i) if the formin can rotate freely around its anchoring point, thermal fluctuations should cause the filament to twirl around its axis, and the polarization index is expected to vary rapidly and erratically over time; (ii) if the formin cannot rotate, then the filament should rotate steadily around its axis as it elongates, and the polarization index is expected to oscillate at a rate matching the filament helicity (~14 full turns per micrometer, at 1 µM actin). The polarization traces were processed by Fast Fourier Transform in order to determine the main frequency of the signal. Filaments exhibiting an erratic rotation (i) were expected to have randomly distributed main frequencies, while filaments exhibiting a steady rotation (ii) were expected to have a main frequency imposed by the filament’s helicity and elongation rate, thus following a normal distribution. Consistently, for each type of formin anchoring, the cumulative distributions of the main frequencies could be well fitted by the weighted sum of a random distribution and a normal distribution (figure 1E). The relative weight of these two distributions thus allowed us to quantify the fraction of filaments in each category. Surprisingly, regardless of the solid (glass) or fluid (lipids) nature of the surface, we found that approximately 54% of filaments elongated by FH1-anchored formins were in situation (i) while this proportion was only 13% for FH2-anchored formins elongated filaments (figure 1F). Therefore, as could be anticipated from the aforementioned structural details, most FH1-anchored formins were able to rotate around their anchoring point, while the rotation of FH2-anchored formins was mostly blocked.

Our results thus far show that the anchored formin’s ability to diffuse on the surface (lipid-versus glass-anchored) and its ability to rotate around its anchoring point (FH1-versus FH2-anchoring) do not significantly affect its ability to rapidly elongate individual actin filaments and to remain attached to their barbed ends.

### Unanchored formins are slower and less processive when filaments are bundled by fascin

We next sought to investigate how filament crosslinking on its own affected the elongation by unanchored formins. We used fascin as a way to crosslink filaments into parallel bundles (figure 2A).

First, we characterized the bundling activity of fascin, using fluorescently labeled fascin. We observed that fascin binds cooperatively to zipper two parallel actin filaments with an effective affinity constant of Kd ~ 75 nM (supp. figure 2A), in agreement with previous reports (Breitsprecher et al., 2011; Winkelman et al., 2014). After two filaments were bundled together by exposure to 100 nM fascin, the bundle could be maintained by fascin concentrations as low as 10 nM, for durations exceeding tens of seconds, in a fascin concentration dependent manner (supp figure 2B, C). This result indicates that bundle maintenance is easier to achieve than its formation. On single filaments (not bundled), the presence of fascin in solution had no detectable impact on formin elongation rate or processivity, for fascin concentrations as high as 1 µM (supp figure 2D).

When filaments elongated by unanchored formins were bundled together by 100 nM fascin, we observed that the barbed end elongation of the leading filament was unaffected, consistent with single filament observations. In contrast, the elongation rates of the first and second trailing filaments were reduced by 18% and 30%, respectively (figure 2B, C, supp figure 2E). This decrease in elongation rate upon fascin-induced bundling is not specific to formins, since we also observed a similar reduction for trailing free barbed ends in bundles formed in the absence of formins (supp figure 2F), as previously reported (Winkelman et al., 2014). This effect could originate from monomer diffusion being partially hindered by the presence of neighboring filaments within the bundle, 6 nm away (Jansen et al., 2011; Yang et al., 2013). One alternative explanation is that the change in filament conformation for trailing filaments upon fascin bundling (Shin et al., 2009) propagates up to the barbed end which would induce a decrease of the affinity for monomeric actin.

We also investigated the impact of filament bundling on the depolymerization of formin-associated barbed ends (figure 2F, G). In the presence of 4 µM profilin and 100 nM fascin, the depolymerization of the leading barbed end was similar to that of single filaments. In contrast, the depolymerization of the trailing barbed end was reduced by 78%, which is a much stronger effect than in the polymerization regime (figure 2C). Remarkably, when the leading formin caught up with the trailing formin, both filament barbed ends depolymerized synchronously, at an intermediate rate, and remained in close proximity (figure 2G). The observed slower trailing formin depolymerization may originate from the combined effect of barbed end conformational change already effective in the polymerization regime, and the stabilization of terminal actin subunits by fascin connections to neighboring filaments. A similar decrease in depolymerization rate and the ensuing barbed end synchronization were observed in the absence of formins (supp. figure 2G).

We next quantified the impact of bundling on the processivity of unanchored formins. In the presence of 100 nM fascin and various concentrations of profilin-actin, the detachment rate of the first and second trailing formins were increased ~3- and 12-fold, respectively, relative to the leading formin, which is unaffected by fascin bundling (figure 2D, supp figure 2H).

To further quantify the impact of bundling on formin activity, we varied the fascin concentration between 50 and 500 nM. At 50 nM fascin, although the filaments visually appeared to be bundled together and trailing formin elongation rates were reduced compared to those of the leading one (supp. figure 2I), we found that, surprisingly, the detachment rate of the leading, first and second trailing formins were all identical (figure 2E). In the presence of 500 nM fascin, the detachment rates of trailing formins were increased to a similar extent as what is observed with 100 nM fascin (figure 2E). Therefore, it appears that low fascin concentrations affect the elongation rate but not the processivity of the formins in the bundle. Overall, these results indicate that formin elongation rate is affected by fascin bundling either through filament overtwisting (Shin et al., 2009) or monomer diffusion hindrance, which happens even at low fascin concentrations, whereas formin processivity appears to be more sensitive to the efficient barbed end zippering by fascin.

### Anchoring formins to a solid surface further hinders their activity in filament bundles

We next sought to investigate how formin anchoring may affect their activity when elongating filaments’ rotation is impaired by crosslinking. In the presence of profilin-actin and 100 nM fascin, as filaments elongate from glass-anchored formins, pairwise bundles form with random distances between the two formins, typically 1 - 20 µm (figure 3A and supp. movie 1). As a standard condition, we used low actin and profilin concentrations to elongate filaments from formins at a moderate rate (5-10 subunits/s) with the intent to keep filaments reasonably short (~10-30 µm), in order to form mainly 2-filament bundles, but similar results were obtained for higher actin and profilin concentrations (see below, figure 3K).

When bundling occurred, we observed that the elongation rates of both leading and trailing formins abruptly changed, dropping on average to 50% of the elongation rate before bundling. In bundles, both the FH1-anchored-to-glass and the FH2-anchored-to-glass formin elongation rates distributed quite uniformly between almost zero and the elongation rate observed before bundling (figure 3B, E). As filament elongation proceeded, two typical behaviors were observed (figure 3G). Either the leading filament elongated faster than the trailing filament and buckled (35%, n=8 out of 23), or the two filaments elongated synchronously (65%, n=15 out of 23). In cases where the leading filament buckled between the leading and trailing formin attachment points, the buckling force was estimated to be lower than 0.15 pN (see Methods). For trailing formins, this force adds up to the minimal viscous pulling force applied to the trailing filament, and is expected to accelerate elongation, not slow it down (Jégou et al., 2013; Kubota et al., 2017; Yu et al., 2017). We therefore concluded that the applied forces are negligible and likely play no role in the slowing down of formin elongation rates.

In these bundles, we also quantified the rate at which filaments detached from the glass-anchored formins. We found that fascin-induced bundling strongly accelerated the detachment of formins (figure 3C, F). The detachment rate in bundles was significantly higher for FH2-anchored-to-glass formins than for FH1-anchored-to-glass formins. Notably, we observed that the processivity of the leading formin was affected by fascin bundling (figure 3H). Leading formins dissociated from barbed ends before trailing formins in 35% of the cases, therefore more frequently than what we observed for unanchored formins (25%). This frequency was independent of the formin anchoring side (figure 3H), of fascin concentration (supp figure 3A, B) and of whether leading filaments buckled or not (supp figure 3C).

Bundled formin detachment rate was independent of formin elongation rate reduction induced by bundling (figure 3J) and of the distance between the leading and trailing formins (when comparing two subpopulations of bundles, with average distances of 2.7 ± 0.8 µm or 7.8 ± 3.3 µm between bundled formins) (figure 3I). When varying profilin-actin concentrations, we observed that bundle formin detachment rates scaled with the single formin detachment rates (figure 3K). We also checked that applying a significant pulling force (by working with stronger microfluidics flow rates) increased formin bundle detachment rate, but did not affect the amplitude of the increase due to bundling (supp figure 3D). Formin detachment rate increase caused by fascin bundling was significantly different depending on the anchoring side, thus on the anchoring details. To further investigate this aspect, we performed measurements with heterodimeric formin constructs, comprising the FH2 dimer and only one FH1 domain and only one His-tag (located either on the FH1-side or the FH2-side, as for homodimers). Each of these formin heterodimers could thus bind to the glass surface via a single anchoring point. Their processivity was reduced to the same extent as for homodimeric formins (supp figure 3E), further validating that the observed formin processivity reduction was not caused by a double anchoring of the formins which would impede their rotation as filaments elongate.

Overall, compared to single formins, the detachment rate of FH1-anchored-to-glass formins was increased 6- and 11-fold, for leading and trailing formins respectively, while for FH2-anchored-to-glass formins this reduction was more pronounced, respectively by 13- and 24-fold (figure 3H). Taken together, those results indicate that, in the situation of bundles where formins are statically anchored to a substrate, the anchoring of formins amplifies the effect of fascin bundling to reduce formin elongation rate and processivity.

### Preventing the rotation of a single filament has the same the impact as bundling on formin activity

To test if the strong reduction in anchored formin activity requires the bundling of filaments by fascin, or if it can occur by only blocking filament rotation, we performed additional experiments where filaments elongating from glass-anchored formins were bound to inactivated glass-anchored NEM-myosins to prevent filament rotation (figure 4A and supp. movie 2). Single formin elongation rates dropped and spread similarly to what we observed for glass-anchored formin bundled by fascin (figure 4B). This observation is reminiscent of what has been observed by Mizuno and colleagues with formin mDia1 elongated filaments occasionally attaching to the surface through biotin-avidin interaction (Mizuno et al., 2018). Formin detachment rate increased upon NEM-myosin binding, similarly to what was observed for leading glass-anchored formins in fascin bundles, with FH2-anchored-to-glass formins detaching faster than FH1-anchored-to-glass formins (figure 4C, D). Using this assay, we thus showed that a single attachment point along the side of the filament is enough to account for most of the change in formin activity observed in fascin bundle experiments. This suggests that the dominant effect of fascin-induced bundling, regarding the activity of glass-anchored formin, is to block filament rotation.

### Lipid-anchored formins efficiently elongate actin filament bundles

In cells, formins are anchored to lipid membranes and can potentially move independently from each other (Higashida et al., 2004). We thus investigated the impact of filament bundling in the more physiological situation where formins are anchored to a lipid bilayer.

Filaments gathered at the edge of the lipid bilayer (figure 5A and supp. movie 3) were bundled together by flowing in fascin, in addition to profilin-actin (figure 5B). Starting with a low density of single filaments allowed us to form mostly 2-filament bundles, whose elongation was then monitored. Here, the distance between bundled formins was much smaller than in the case of glass anchored formins, and below the optical resolution of our TIRF microscope (~200 nm). Therefore we used fluorescent fascin to monitor when the bundled region, labeled by fluorescent fascin, moved away from the lipid edge (figure 5C), which we identified as formin dissociation events. 2-filament bundles grew steadily and the elongation rate was not reduced compared to single filaments (figure 5D). For some bundles, we could observe leading formins moving upstream as they elongated filament barbed ends, away from the lipid pattern edge and the trailing formin (figure 5C, bottom), with a difference in elongation rate between leading and trailing formins similar to what we observed for unanchored formins.

For 2-filament bundles, the detachment rate of lipid-anchored formins appeared similar to that of unanchored formins (figure 5F). The ~2-fold difference between FH1-anchored-to-lipids and FH2-anchored-to-lipids formins detachment rate observed for glass-anchored formins was still present for lipid-anchored formins, indicating that the tether length still plays an important role even for lipid-anchored formins in a bundle.

Overall, the results obtained from lipid-anchored formins show that (1) the translational freedom of formins thanks to the fluidity of the lipid bilayer seems to allow a smoother and more processive filament elongation by formins when filament rotation is blocked by fascin bundling, while (2) this process is still dependent on the tether length connecting the formins to the surface.

## Discussion

In this work, we investigated the impact of geometrical constraints on the activity of formin mDia1, quantified by the rate at which it elongates barbed ends and by the rate at which it detaches from them, which we summarized figure 6. We found that the application of constraints to both actin filaments (bundling) and formins (anchoring) had a significant impact on formin activity (figure 3), while these constraints taken separately had little or no effect (figures 1 and 2). This unsuspected cross-talk between constraints located micrometers apart is illustrated by our observation that formin anchoring details, which appear to play no role when elongating individual filaments, matter significantly as soon as filaments are bundled. In particular, constraints relative to formin’s ability to rotate around its anchoring point appear to have a strong impact on formin activity when filaments are bundled. Remarkably, we found that formins anchored to a lipid bilayer via their N-terminus, hence with minimal constraints, are able to elongate bundled filaments as efficiently as if they were not anchored (figure 5).

**Figure 6:**
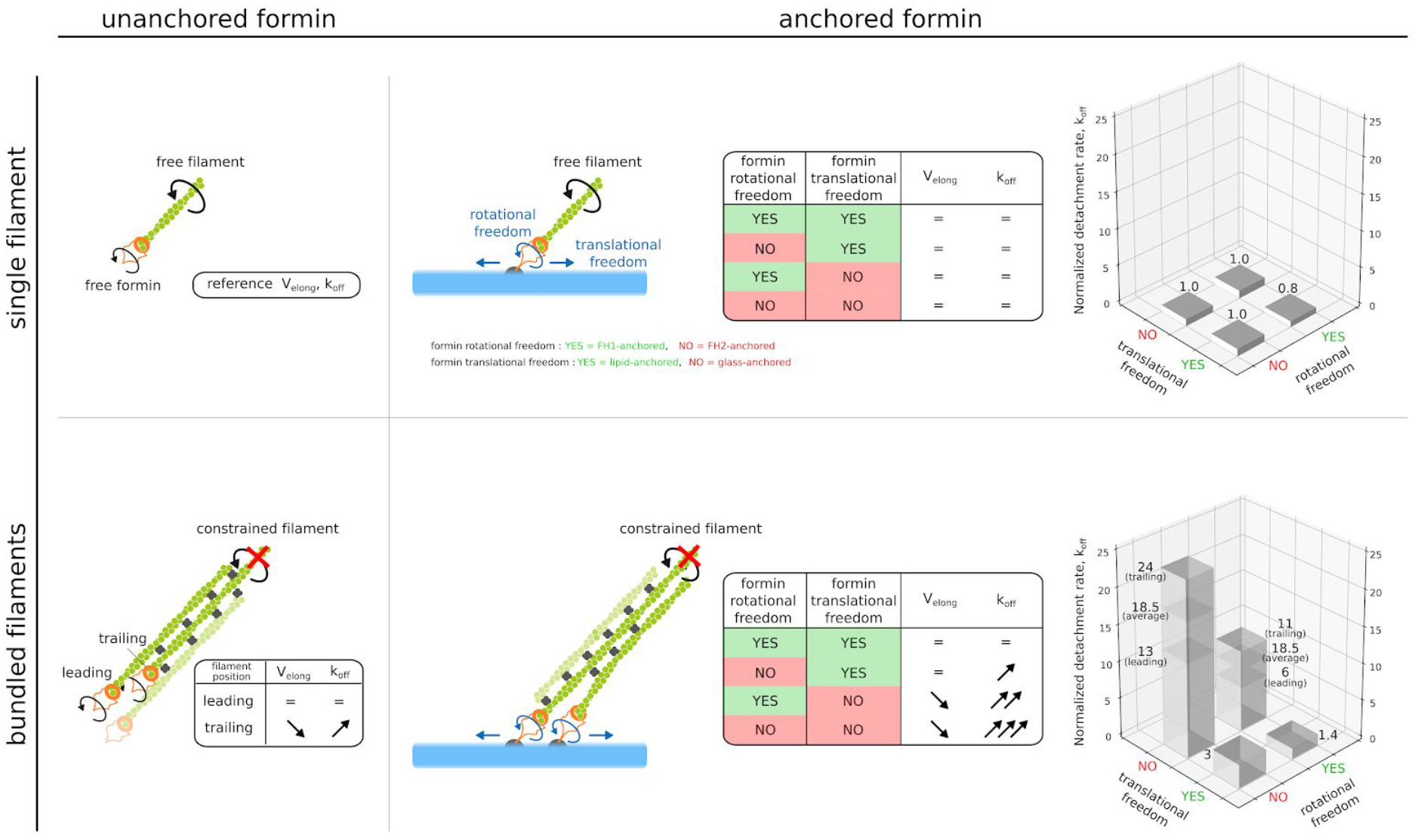
Schematic summary of the impact of filament bundling and formin anchoring on formin activity. Depending on filaments being bundled by fascin or not, and whether formin translational or rotational freedom are blocked or not, formin-induced actin filament elongation and formin detachment rate are differently affected. Top, left: a single filament elongated by an unanchored formin is chosen as the reference for the comparison of elongation rates (V_elong_) and formin detachment rates (k_off_). Bottom, left: when fascin bundles filaments, the activity of the formin elongating the leading filament is unaffected, while the elongation rate is reduced and the detachment rate is increased for formins elongating trailing filaments. Top, right: the activity of surface-anchored formins elongating individual filaments is not affected by anchoring. Bottom, right: Simultaneously blocking filament rotation (for example by fascin bundling) and formin rotational or translational freedom, impacts strongly both formin elongation rate and detachment rate for both leading and trailing filaments.

Interestingly, the issue of formin’s rotational freedom upon anchoring has recently been investigated in the context of pulling forces applied to single filaments by Yu et al. (Yu et al., 2017). In this study, the authors used magnetic beads to pull on filaments and found that piconewton pulling forces applied to formins that were free to rotate allowed them to elongate filaments extremely rapidly, close to the diffusion limit (Yu et al., 2018). In contrast, when applying significant forces to our freely rotating formins, the elongation rates remained comparable to what we have reported earlier using microfluidics (Cao et al., 2018; Jégou et al., 2013), and to what others have reported using optical traps (Kubota et al., 2017) or myosins (Zimmermann et al., 2017) to apply pulling forces. Control experiments in the latter study indicate that the effect they observe (on mDia2 and Cdc12 formins) can indeed be attributed to tension. Nonetheless, the contribution of active myosins to torsional constraints is not fully understood (Leijnse et al., 2015) and may be worth investigating further, following our results.

Thanks to these earlier reports that used different means to apply mechanical tension to actin filaments, the sensitivity of formins to external forces is now well established (Zimmermann and Kovar, 2019). Here, we show that formins are also sensitive to the geometrical organization of the filaments they elongate. When filaments are crosslinked and cannot rotate around their main axis, hindering the rotation of the anchored formin results in the generation of a mechanical torque as the filaments elongate, and as a consequence the formin’s elongation rate is reduced and its off-rate is increased. This sensitivity to geometry, where mechanical stress is not applied from the outside but rather generated by the protein’s activity in a specific context, is reminiscent of our recent work showing that cofilin binding applies a mechanical torque to cross-linked filaments, and thereby greatly enhances their severing (Wioland et al., 2019).

Here, we focused on fascin-induced bundling as a means to cross-link filaments, a situation typically encountered in filopodia (Vignjevic et al., 2006) and invadopodia (Li et al., 2010). More generally, most actin crosslinkers and geometries are likely to also block filament rotation and to have a similar impact on formin activity, as exemplified by our experiments with NEM-myosins (figure 4). As we show, the details of formin anchoring are of great importance in this context, and these can certainly take different forms in cells. The binding of Diaphanous related formins directly to membranes or to membrane-bound proteins and activators via their N-terminal regions (Aspenström, 2010; Ramalingam et al., 2010; Rousso et al., 2013) likely allows them to maintain some rotational freedom while being anchored. This freedom may be tuned by a number of other factors, such as the lipid composition of the membrane or the presence of other proteins. In particular, in the context of filopodia, the presence of Ena/VASP and its potential interaction with formins (Barzik et al., 2014; Bilancia et al., 2014), may alter formin’s rotational freedom and thereby its ability to efficiently elongate barbed ends for significant durations. Future studies will certainly shed light on how the rotational freedom of membrane-anchored formins can be modulated and regulate the assembly dynamics of actin filaments in cells.

## Acknowledgements

We thank all members of the G.R.-L./A.J. laboratory for their help, and especially Hugo Wioland for help in the fluorescence polarization analysis and Sandy Jouet for protein purification. We acknowledge funding from the French ANR (Grants ‘MuscActin’ and ‘Conformin’ to G.R.-L.) and the European Research Council (Grant StG-679116 to A.J.).

